# Gene expression markers of Tumor Infiltrating Leukocytes

**DOI:** 10.1101/068940

**Authors:** Patrick Danaher, Sarah Warren, Lucas Dennis, Leonard D’Amico, Andrew White, Mary L. Disis, Melissa A. Geller, Kunle Odunsi, Joseph Beechem, Steven P. Fling

## Abstract

**Background:** Assays of the abundance of immune cell populations in the tumor microenvironment promise to inform immune oncology research and the choice of immunotherapy for individual patients. We propose to measure the intratumoral abundance of various immune cells populations with gene expression. In contrast to IHC and flow cytometry, gene expression assays yield high information content from a clinically practical workflow. Previous studies of gene expression in purified immune cells have reported hundreds of genes showing enrichment in a single cell type, but the utility of these genes in tumor samples is unknown. We describe a novel statistical method for using co-expression patterns in large tumor gene expression datasets to validate previously reported candidate cell type marker genes, and we use this method to winnow previously published gene lists down to a subset of high confidence marker genes.

**Methods:** We use co-expression patterns in 9986 samples from The Cancer Genome Atlas (TCGA) to validate previously reported cell type marker genes. We compare immune cell scores derived from these genes to measurements from flow cytometry and immunohistochemistry. We characterize the reproducibility of our cell scores in replicate runs of RNA extracted from FFPE tumor tissue.

**Results:** We identify a list of 60 marker genes whose expression levels quantify 14 immune cell populations. Cell type scores calculated from these genes are concordant with flow cytometry and IHC readings, show high reproducibility in replicate RNA samples from FFPE tissue, and reveal an intricate picture of the immune infiltrate in TCGA. Most genes previously reported to be enriched in a single cell type have co-expression patterns inconsistent with cell type specificity.

**Conclusions:** Due to their concise gene set, computational simplicity and utility in tumor samples, these cell type gene signatures may be useful in future discovery research and clinical trials to understand how tumors and therapeutic intervention shape the immune response.

## Background

The abundance and composition of the immune cells infiltrating a tumor predict both a patient’s prognosis [1-5] and the optimal immunotherapy for their disease [6]. Therefore techniques to profile tumor infiltrating lymphocytes (TILs) in a clinical setting are needed. Flow cytometry provides a gold standard for quantifying immune cell populations, but its complex workflow, expense, need for high numbers of cells, and long processing time make it unavailable to most clinics. Immunohistochemistry (IHC) has been shown to be clinically useful [2], but it cannot assay more than a few immune markers without using up excessive tissue. In contrast to these older technologies, gene expression profiling promises a clinically practical way to measure the full diversity of the tumor immune infiltrate, requiring limited tissue and allowing the simultaneous measurement of hundreds to thousands of clinically relevant genes. We propose to identify genes whose expression levels can be used to measure the abundance of various immune cell populations within the tumor microenvironment.

Previous authors have identified genes specific to purified immune cell populations [7-9] and used these genes to quantify immune populations in tumors [9,10]. However, these genes were discovered using purified cells and not immune cells taken from the tumor microenvironment, and so any differences between intratumoral and in vitro gene expression patterns will introduce noise into their measurements. To address this concern, we propose a novel computational method for testing whether previously reported cell type marker genes are useful in tumor data. We then apply this method to data from The Cancer Genome Atlas (TCGA) to derive a set of 60 validated marker genes for 14 immune cell populations.

Our final gene list exhibits sufficiently strong cell type specificity to allow measurement of immune cell populations with scores computed as the simple average log expression of their marker genes. In data from ovarian cancer patients, these cell type scores show strong concordance with both immunohistochemistry (IHC) and flow cytometry. In replicate RNA samples, they display substantially better reproducibility than typical IHC readings. In TCGA data, they reveal an intricate picture of anti-tumoral immunity. Due to their concise gene set, computational simplicity and utility in tumor samples, these cell type gene signatures may be useful in future discovery research and clinical trials to understand how tumors and therapeutic intervention shape the immune response.

## Methods

### Pairwise similarity statistic for quantifying marker-like co-expression patterns

If two genes are ideal cell type markers, their log expression values will be perfectly correlated with a slope of 1. The below adaptation of Pearson’s correlation metric measures a pair of genes’ adherence to this pattern:

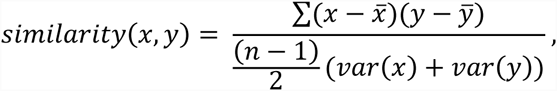

where x and y are the vectors of log-transformed, normalized expression values of the two genes, 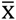 and 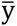 are their sample means, and var(x) and var(y) are their sample variances. This function equals 1 when the two genes are perfectly correlated with a slope of 1 and decreases for gene pairs with low correlation or with slope diverging from 1. Since many biologically related genes will exhibit correlation unrelated to a shared cell type, mere correlation is a weak indicator of cell type markers. Similarly, gene pairs that exhibit pairwise differences with low variance are consistent with the hypothesis that they serve as cell type markers, but unless they retain this stable pairwise difference over a range of expression values and thereby achieve high correlation, they provide minimal evidence for their utility as cell type markers.

### Flow cytometry

Whole blood was stained within 30 hours of collection with 2 12-color antibody staining panels: a PBMC subset panel and a T cell subset panel. The PBMC subset panel antibody cocktail (CD3-PE CF594, CD4-FITC, CD8-PerCP Cy5.5, CD11c-AlexaFluor 700, CD14-V450, CD16-APC H7, CD19-PE-Cy5, CD45-AmCyan, CD56-PE Cy7, CD122 APC, CD123 PE, HLADR-BV605) was used to stain 100uL whole blood in BD Trucount Tubes to determine absolute numbers of various peripheral immune cell types, including monocytes, CD4 and CD8 T cells, NK cells, NKT cells, B cells, plasmacytoid dendritic cells (pDC) and myeloid dendritic cells (mDC). After staining, cells were incubated with FACs Lysing Solution (BD) for 15 minutes, and stored at -80°C until acquisition. For determining activation state, as well as naïve-memory-effector subsets of CD4 and CD8 T cells, 200 uL of whole blood was incubated with the T cell subset panel antibody cocktail (CD3-FITC, CD4-APC-Cy7, CD8-PerCP Cy5.5, CD25-APC, CD28-PE-CF594, CD45-AmCyan, CD45RA-BV650, CD127-BV421, CD197-AlexaFluor 700, CD278-PE, CD279-PE-Cy7, HLADR-BV605), and subsequently lysed with Pharm Lyse solution (BD), washed, fixed with 1% PFA, and suspended in 10% DMSO before storage at -80°C. Samples from both panels were acquired on a BD LSR II cell analysis machine and analyzed by FlowJo Cell Analysis software.

### TCGA data

Normalized RNASeq data was downloaded from TCGA and log2-transformed prior to analysis. No further preprocessing was applied.

### NanoString data

RNA from PBMC lysates (~60,000 cells per assay) and FFPE tumor biopsy sections (150-300ng per assay) were evaluated for gene expression using the nCounter PanCancer Immune Profiling panel, which interrogates 770 immune-related genes and associated controls.

NanoString gene expression values were normalized using the best subset of the 40 reference genes included in the panel, as determined by geNorm [11].

### Statistical Methods: Comparison to flow cytometry and IHC

We measured concordance between platforms with Pearson correlation and Root Mean Squared Error (RMSE). RMSE between matching measurements from NanoString and either flow or IHC was calculated by mean-centering each separate set of measurements and then taking the square root of the mean squared difference between matching pairs.

### Statistical Methods: Reproducibility analysis

To measure the proportion of variance due to noise for each cell score, we fit a linear mixed model predicting cell score from sample ID, treating sample ID as a random effect. As a measure of the proportion of variance due to noise, we report the estimated residual variance divided by the sum of the residual variance and the between-sample variance.

## Results

### Derivation of literature-derived candidate cell type marker genes

The first step in our marker gene identification process is to identify previously reported cell type markers. Fortunately, the literature is rich in papers measuring gene expression in isolated immune cell populations. A number of authors, most notably [7] and [9], have used meta-analyses of these experiments to discover genes that are predominantly expressed within a single immune cell population. Our list of candidate marker genes is primarily drawn from [7], using [9]’s list to fill in cell types absent from [7]. The list also includes well-known markers for exhausted CD8 cells [12-14] and FOXP3 for Tregs.

### Approach to validation of candidate cell type marker genes in tumor gene expression data

The literature on cell type specific gene expression is a powerful source of candidate marker genes, but there are a variety of mechanisms by which poor marker genes may have entered the literature. First, early microarray studies were frequently underpowered, noisy and rife with batch effects and therefore could indicate spurious marker genes [15]. Second, and very significantly, the expression profile of in vitro purified cells may differ substantially from these cells’ gene expression in the tumor microenvironment [16]. Finally, genes that appear specific to one cell type in a microarray experiment may be expressed in cell types omitted from the experiment.

Therefore, our literature-derived candidate marker genes require validation in actual tumor expression data. Ideally, we would test whether each gene displays two properties: expression specific to a single cell type, and stable expression within that cell type. Unfortunately, we cannot directly measure a gene’s adherence to these properties in bulk tumor expression data; instead, we look for genes whose expression patterns are consistent with these properties. If two genes are both ideal markers, expressed with perfect specificity to and stability within a cell type, their expression levels will be perfectly correlated, and the ratio between them will be constant across samples. Figure 1 demonstrates this principle. Of the 4 candidate marker genes for a cell population, Genes 1 and 2 rise and fall at the same rate, a co-expression pattern consistent with both genes being driven by abundance of a single cell population. By contrast Gene 3 exhibits no such co-expression with Gene 1 and likely does not serve as a marker gene in the tumor microenvironment. Gene 4 is highly correlated with Genes 1 and 2, but its slope is different than 1. Thus while Genes 1, 2 and 4 may be regulated by the same biological process, that they do not increase at the same rate means they are not all expressed at consistent levels within a single cell type. We quantify genes’ adherence to the marker-like co-expression pattern we seek with a pairwise similarity statistic, defined in the Methods.

**Figure 1:**
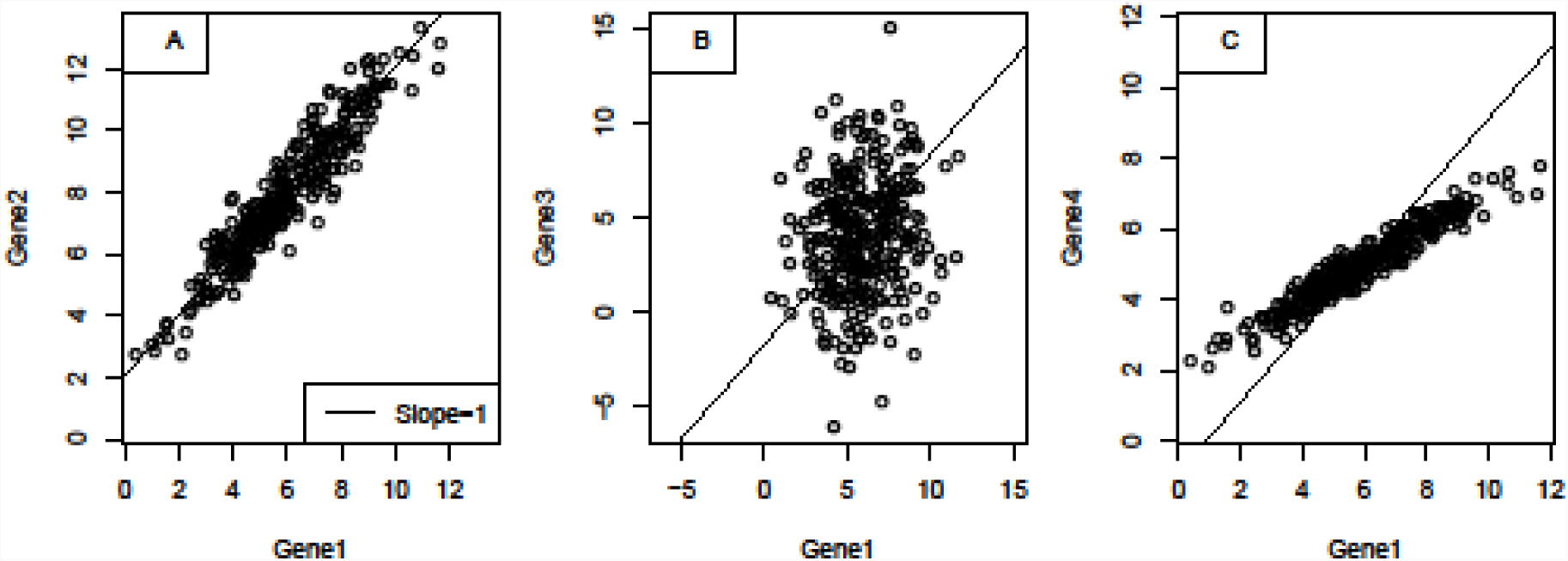
Simulated example of process for evaluating candidate genes for marker-like co-expression in a tumor dataset. A) Genes 1 and 2 are highly correlated with a slope of 1, a pattern consistent with both genes rising and falling at the same rate. B) Gene 3 exhibits no such co-expression with gene 1, showing that Genes 1 and 3 are not both good markers for the same cell type. C) Gene 4 is highly correlated with Gene 1, but with a slope different than 1, meaning they are not both markers for the same cell type.

For each set of candidate marker genes for a single cell type, we sought a subset of genes that exhibited strong marker-like co-expression patterns in tumor gene expression data. We considered candidate marker genes with expression patterns like Genes 1 and 2 to be validated, and we discarded candidate markers that behaved like Gene 3 or Gene 4. At all times we allowed well-established biology to inform the selection process. For example, we discarded a cluster of putative Th2 cell marker genes (BIRC5, HELLS, CDC7, WDHD1, CENPF, NEIL3, DHFR, DC25C) that showed strong marker-like co-expression (Supplementary Figures) but had many genes that were previously reported to be expressed broadly across cell types [17-19].

Alone, co-expression patterns are insufficient to establish a group of genes as markers for a single cell type. However, when a set of genes has been previously reported to have cell type-specific expression and also displays marker-like co-expression patterns in tumor data, it provides strong evidence for their use as cell type markers.

### Using marker genes to quantify cell types

Once we have selected a set of marker genes for a cell type, measuring the cell type’s abundance is straightforward. Assuming each marker gene is present at a fixed but unknown number of copies per cell, the average log-transformed expression of the marker genes is equal to the log-transformed abundance of the cell type, plus an unknown constant. Thus we compute cell type scores with the simple average of their marker genes’ log-transformed expression values. Because of the unknown constant, these scores do not provide absolute quantification of cell types; e.g., we cannot say, ^“^there are 500 CD8 cells in this sample,^”^ or, ^“^this sample has more B-cells than T-cells.^”^ But they do allow comparison of cell abundance across samples, e.g., ^“^this tumor has twice as many CD8 cells as the average tumor,^”^ sufficient information for many scientific and clinical applications. If our scores are calculated from log2 transformed data, each unit increase in a cell score should correspond to a doubling of that cell type’s abundance.

### Only a small proportion of previously reported cell type specific genes display marker-like co-expression patterns in the tumor microenvironment

The Cancer Genome Atlas (TCGA) provides ideal data for validating candidate cell type marker genes through their co-expression patterns. We evaluated our literature-derived candidate marker genes in TCGA RNASeq data from 9986 samples from 32 tumor types. Details of the TCGA download used are in Table S1.

TCGA data revealed previously reported cell type marker genes to have widely varying quality, with many candidate cell type marker genes displaying co-expression patterns inconsistent with cell type specificity and stability. However, most cell types had a core subset of genes with strong marker-like co-expression (Figures 2, 3, Supplementary Figures). These highly concordant gene sets constitute our final selected cell type markers (Table 1).

**Figure 2:**
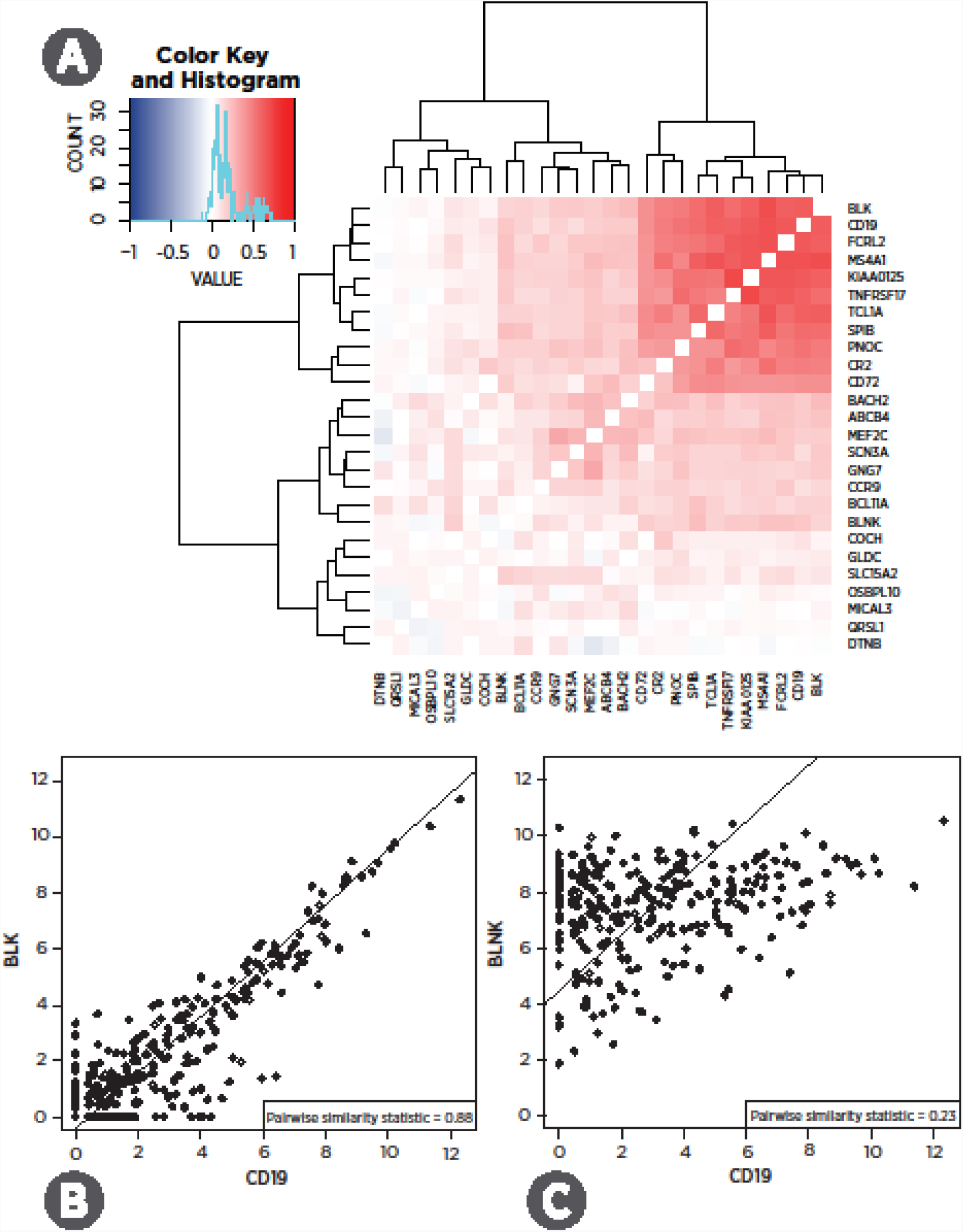
Pairwise similarity, a measure of marker-like co-expression, of candidate B-cell marker genes in TCGA. a) Pairwise similarity of candidate B-cell marker genes averaged across 24 TCGA RNASeq datasets. Darker red indicates co-expression patterns consistent with both genes acting as cell type markers. Values of 1indicate perfect marker-like co-expression. Green sidebars indicate final selected markers. b) Two of the selected B-cell markers, including CD19, in the bladder cancer dataset. c) In bladder cancer, CD19 and a candidate B-cell marker that we discarded.

**Figure 3:**
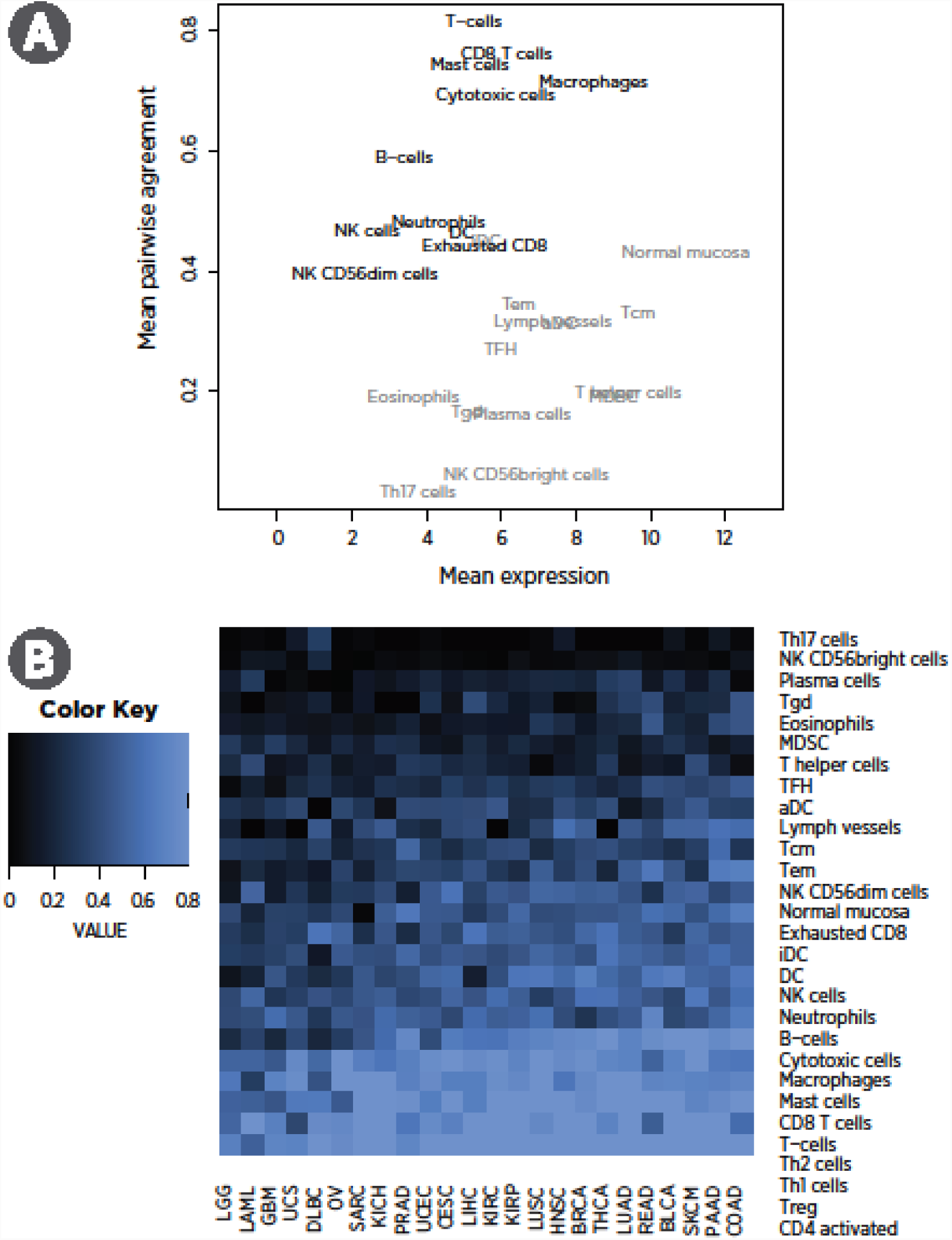
Pairwise similarity, a measure of marker-like co-expression, of candidate marker genes in TCGA. a) Mean log2 expression vs. average pairwise similarity of selected cell type markers across TCGA datasets. Cell types in grey have been discarded from the final panel of markers. b) Average pairwise similarity of each cell type’s marker genes in each TCGA dataset.

**Table 1:** Summary of the marker gene selection results for each cell type. Cell types lacking validated marker genes are omitted. The mean pairwise similarity statistic is a measurement of how well a gene set adheres to the co-expression patterns expected from a set of perfect marker genes, with a score of 1 indicating perfect marker-like behavior.

| Cell type | # Candidate genes | # Selected markers | Mean pairwise similarity statistic in TCGA | Selected marker genes |
| --- | --- | --- | --- | --- |
| B-cells | 34 | 9 | 0.59 | BLK, CD19, FCRL2, MS4A1, KIAA0125, TNFRSF17, TCL1A, SPIB, PNOC |
| CD45 | 1 | 1 | *NA | PTRPC |
| Cytotoxic cells | 18 | 10 | 0.69 | PRF1, GZMA, GZMB, NKG7, GZMH, KLRK1, KLRB1, KLRD1, CTSW, GNLY |
| DC | 7 | 3 | 0.46 | CCL13, CD209, HSD11B1 |
| Exhausted CD8 | 5 | 4 | 0.44 | LAG3, CD244, EOMES, PTGER4 |
| Macrophages | 33 | 4 | 0.71 | CD68, CD84, CD163, MS4A4A |
| Mast cells | 31 | 5 | 0.74 | TPSB2, TPSAB1, CPA3, MS4A2, HDC |
| Neutrophils | 32 | 7 | 0.48 | FPR1, SIGLEC5, CSF3R, FCAR, FCGR3B, CEACAM3, S100A12 |
| NK CD56dim cells | 14 | 4 | 0.40 | KIR2DL3, KIR3DL1, KIR3DL2, IL21R |
| NK cells | 36 | 3 | 0.47 | XCL1, XCL2, NCR1 |
| T-cells | 13 | 6 | 0.81 | CD6, CD3D, CD3E, SH2D1A, TRAT1, CD3G |
| Th1 cells | 27 | 1 | *NA | TBX21 |
| Treg | 18 | 2 | *NA | FOXP3 |
| CD8 T cells | 35 | 2 | 0.51 | CD8A, CD8B |
| CD4 cells | 20 | 0 | **NA |  |

\* Single marker gene; quality impossible to assess in expression data alone
\*\* Calculated as the T-cell score minus the CD8 cell score

Figure 2 illustrates our selection process: of the 26 genes previously reported as being expressed specifically in B-cells, most have co-expression patterns incompatible with specificity to the same cell type. But a subset of genes, including the canonical B-cell marker CD19, share the co-expression patterns we seek, namely high correlation with a slope near 1 (Figure 2a). For example, in the TCGA bladder cancer (BLCA) dataset, BLK and CD19 show nearly perfect marker-like co-expression (Figure 2b), while the putative B-cell marker BLNK is largely uncorrelated with CD19 (Figure 2c). BLNK’s unsuitability as a B-cell marker is corroborated by [20]’s finding of BLNK expression in murine macrophages.

Cell types varied in the quality of their selected marker genes (Figure 3). For example, our selected set of T-cell genes showed very strong marker-like co-expression, while our selected T-helper cell genes displayed weak marker-like co-expression (Supplementary Figures). Genes with lower average expression were less likely to display expression patterns typical of ideal cell type markers, a pattern consistent with greater measurement error at low expression values. Noting this pattern, two clusters of cell types with respectively successful and unsuccessful marker genes are apparent (Figure 3a). We discarded the cluster of cell types whose marker genes were unimpressive given their expression level.

In two cases, we retained a single gene as a cell type marker. For Th1 cells, we found no clusters of candidate genes with marker-like co-expression; thus we selected TBX21, the gene for the classic Th1 cell marker TBET. However, [21] reported T-bet expression in B-cells, so this marker gene may be influenced by B-cell abundance. We also use PTRPC as an unvalidated, single-gene marker CD45+ cells, although it is likely expressed at different levels by different cell types.

The selected marker genes appear to have pan-cancer utility: each set of marker genes showed similar performance across TCGA datasets. An important exception is brain and immune tumors, which showed reduced marker-like co-expression for all cell types (Figure 3b). Poor performance in immune tumors might be expected to result from tumor-intrinsic expression of immune genes, and poor performance in brain tumors likely results from the blood-brain barrier limiting the dynamic range of immune cell abundance and thereby limiting our ability to resolve marker-like co-expression patterns.

### Comparison of gene expression cell type scores to flow cytometry and IHC

FFPE tissue and PBMCs were collected from ovarian cancer patients. CD3+ and CD8+ cells were quantified in FFPE samples using IHC, and numerous cell populations (Table S2) were quantified in PMBCs using flow cytometry. In 19 FFPE and 18 PBMC samples, we measured expression levels of our 60 cell type marker genes and of 670 additional genes relevant to the tumor-immune interaction. Gene expression cell type scores were broadly concordant with both flow cytometry and IHC measurements (Figure 4, Table 2).

**Figure 4:**
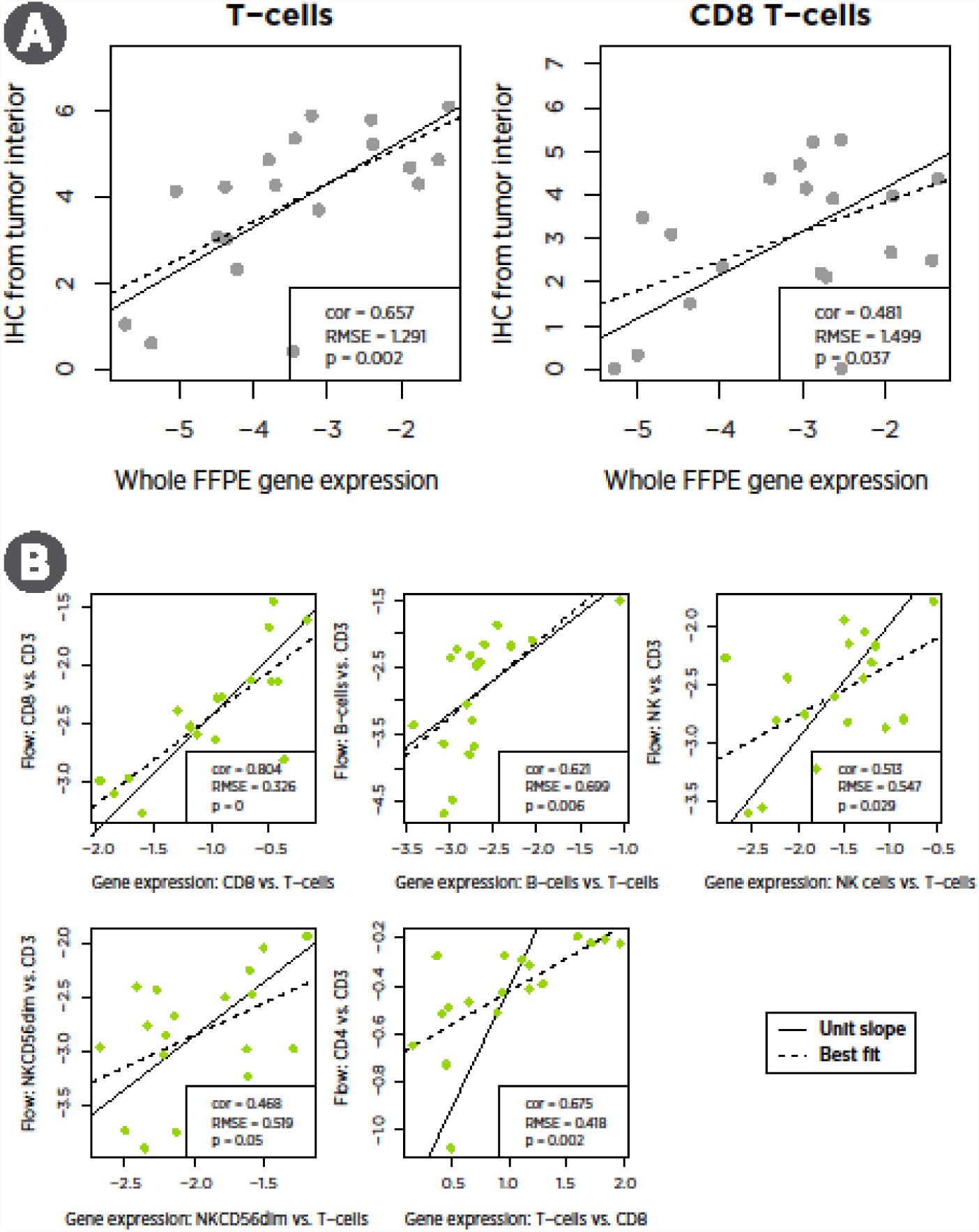
a) Comparison of gene expression and IHC cell type measurements in FFPE tumor samples. b) Comparison of gene expression and flow cytometry cell type measurements in PBMCs.

**Table 2:** Summary of evidence for each cell type score. Root mean squared errors are on the log2 scale. The mean pairwise similarity statistic measures how well a gene set’s co-expression pattern adheres to the co-expression pattern of ideal marker genes, with a value of 1 indicating perfect correlation with a slopeof 1. The standard deviation (SD) and proportion of variance due to noise were calculated from triplicate geneexpression assays from tumor sample RNA.

| Cell type | Correlation with IHC | Root Mean Squared Error from IHC | Correlation with flow | Root Mean Squared Error from Flow | Mean pairwise similarity statistic in TCGA | SD due to technical noise (log2 scale) | Proportion of variance due to noise |
| --- | --- | --- | --- | --- | --- | --- | --- |
| B-cells |  |  | 0.62 | 0.064 | 0.59 | 0.13 | 0.0022 |
| CD45 |  |  |  |  | ***NA | 0.1249 | 0.0024 |
| Cytotoxic cells |  |  |  |  | 0.69 | 0.0813 | 0.001 |
| DC |  |  |  |  | 0.46 | 0.2307 | 0.0151 |
| Exhausted CD8 |  |  |  |  | 0.44 | 0.1624 | 0.0062 |
| Macrophages |  |  |  |  | 0.71 | 0.0828 | 0.0013 |
| Mast cells |  |  |  |  | 0.74 | 0.1949 | 0.0086 |
| Neutrophils |  |  |  |  | 0.48 | 0.19 | 0.0026 |
| NK CD56dim cells |  |  | 0.47 | 0.071 | 0.40 | 0.2347 | 0.1073 |
| NK cells |  |  | 0.51 | 0.118 | 0.47 | 0.1938 | 0.017 |
| T-cells | 0.66 | 1.30 | *0.78 | *0.064 | 0.81 | 0.1116 | 0.0021 |
| Th1 cells |  |  |  |  | ***NA | 0.2212 | 0.0304 |
| Treg |  |  |  |  | ***NA | 0.371 | 0.049 |
| CD8 T cells | 0.53 | 1.50 | 0.78 | 0.138 | 0.51 | 0.1842 | 0.0045 |
| CD4 cells |  |  | **0.65 | **0.752 |  |  |  |

\* Used to normalize the other cell types; 0.78 and 0.064 are the highest correlation and lowest RMSE observed between gene expression and flow for any T-cells vs. other cell type contrast.
\*\* Calculated as the T-cell score minus the CD8 cell score
\*\*\* Only one marker gene; quality impossible to assess in expression data alone

For comparison of our cell scores to flow cytometry, the normalization of gene expression data in PBMCs required a non-standard method. Changes in the composition of PBMCs can influence the abundance of housekeeping/reference genes, spuriously changing normalized expression values and by extension our cell type scores. We avoided this problem by normalizing our cell type scores not to reference genes but to our T-cells score, which appears to be our most accurate score. Several of our cell type scores lacked an exact counterpart in the flow cytometry data and thus could not be validated by this method.

A notable finding from the flow cytometry data is the ability of our cell type scores to predict CD4 abundance. Although we have no explicit CD4 cell score – our analysis of TCGA data cast doubt on the utility of all the reported T helper cell genes – the difference between our T cell and CD8 cell scores correlates strongly (r=0.65) with the difference between flow cytometry CD4 and CD3 log counts.

The between platform correlations in cell type measurements were in general moderately strong but statistically significant. The near-unit slopes of the lines of best fit in these plots is important: these slopes mean that a 2-fold increase in T-cells as measured by gene expression predicts a 2-fold increase in T-cells as measured by IHC. Note that each platform returns results on a different scale, and so it is necessary to mean-center their measurements before comparing them.

The low reproducibility of IHC measurements [22] and the variable spatial distribution of immune cells within a tumor sample place strong upper bounds on the correlation between IHC and gene expression measurements of cell type abundance. For example, spatial sampling factors appear to explain the low outlier in Figure 4: in this IHC sample, CD3 and CD8 cells were nearly absent from the tumor interior but were highly abundant in the invasive margin.

The correlation between gene expression and flow cytometry is limited by the relatively constant proportions of immune cell populations in PBMCs. The most variable comparison, CD4 cells vs. CD3 cells, changed by less than 30% between its minimum and maximum. In contrast, IHC T-cell measurements increased by 20-fold between their minimum and maximum. The very small root mean squared errors (RMSE) between gene expression and flow measurements are consistent with high concordance but low variance. Further discordance between gene expression and flow cytometry can be attributed to measurement errors in both platforms, gating decisions in flow analysis, and genuine differences in the biology captured by the two platforms.

### Reproducibility of gene expression cell type scores

To evaluate our cell scores’ technical reproducibility, we assayed RNA extracted from 12 tumor FFPE samples (Asterand) in triplicate using the nCounter PanCancer Immunology Panel (NanoString Technologies). These 12 samples included 2 endometrial carcinomas, 3 cervical carcinomas, 2 thyroid carcinomas, 2 neuroendocrine carcinomas, 2 esophageal tumors, and 1 mesothelioma. Reproducibility for most cell scores was extremely high (Figure 5, Table 2), with the median cell score having a negligible 0.5% of variance explained by technical noise. Our NK CD56 Dim cell score had notably worse reproducibility than the rest, with 10% of variance due to noise.

**Figure 5:**
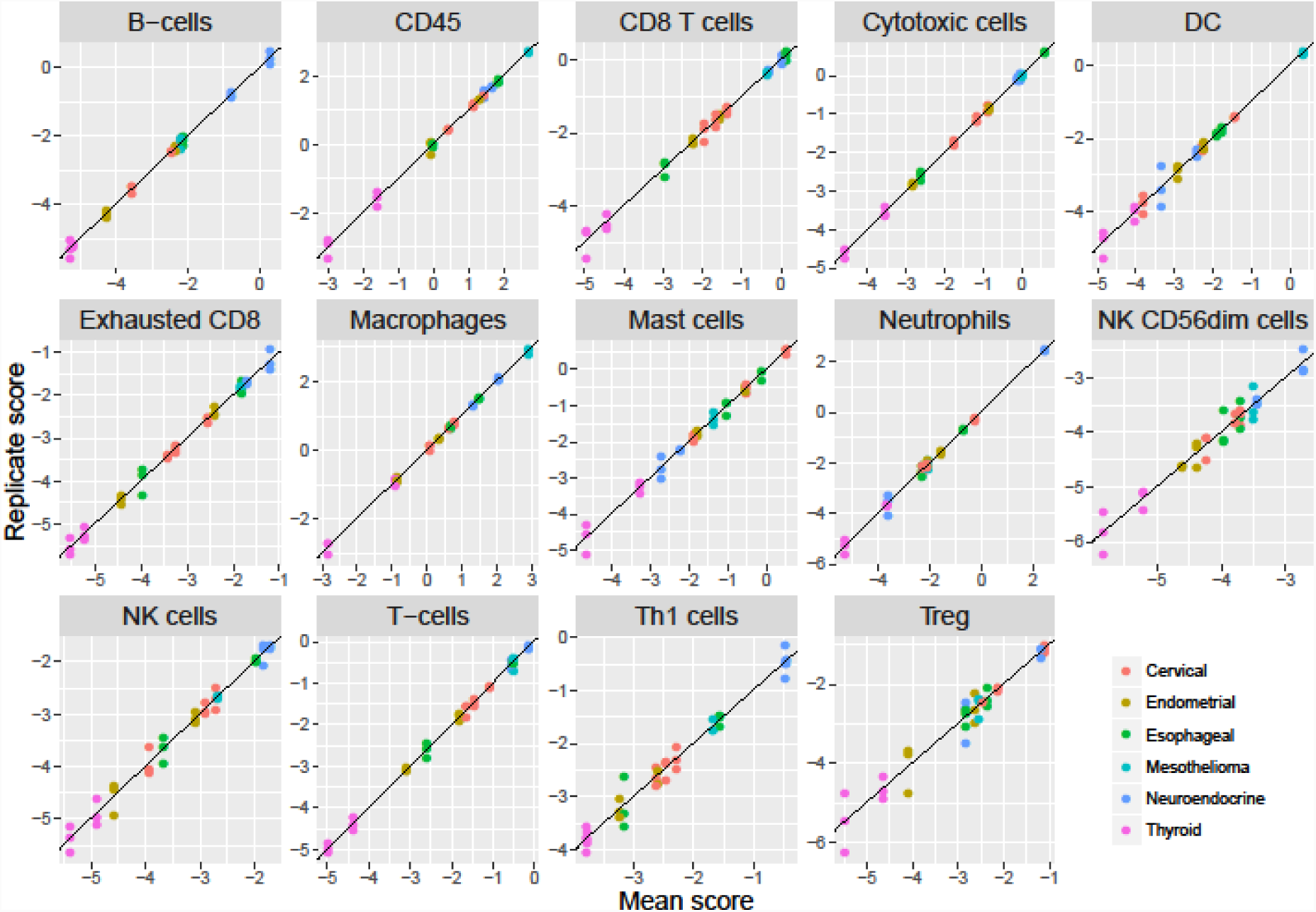
reproducibility of cell scores derived from triplicate runs of 12 tumor samples.

### Application of cell type marker genes to TCGA RNASeq data

#### Results in TCGA: pan-cancer patterns in TIL abundance

We used our immune cell marker genes to calculate cell abundance scores in 9986 TCGA RNASeq samples from 24 tumor types. The majority of immune cell scores tended to rise and fall together, with the average pair of cell scores having a correlation of 0.61 over the solid tumor TCGA datasets. This finding suggests that the primary component of variance in most cell types’ abundance is driven by the amount of infiltrate rather than its makeup. To capture this primary axis of information, we defined a ^“^total TILs^”^ signature as the average of all cell scores with correlations with PTRPC (CD45) greater than 0.6, which excluded only dendritic cells, Tregs and mast cells. Out total TILs score explained 60% of the variance in our cell scores in TCGA data. Total TIL score varied widely between and within tumor types (Figure 6b).

**Figure 6:**
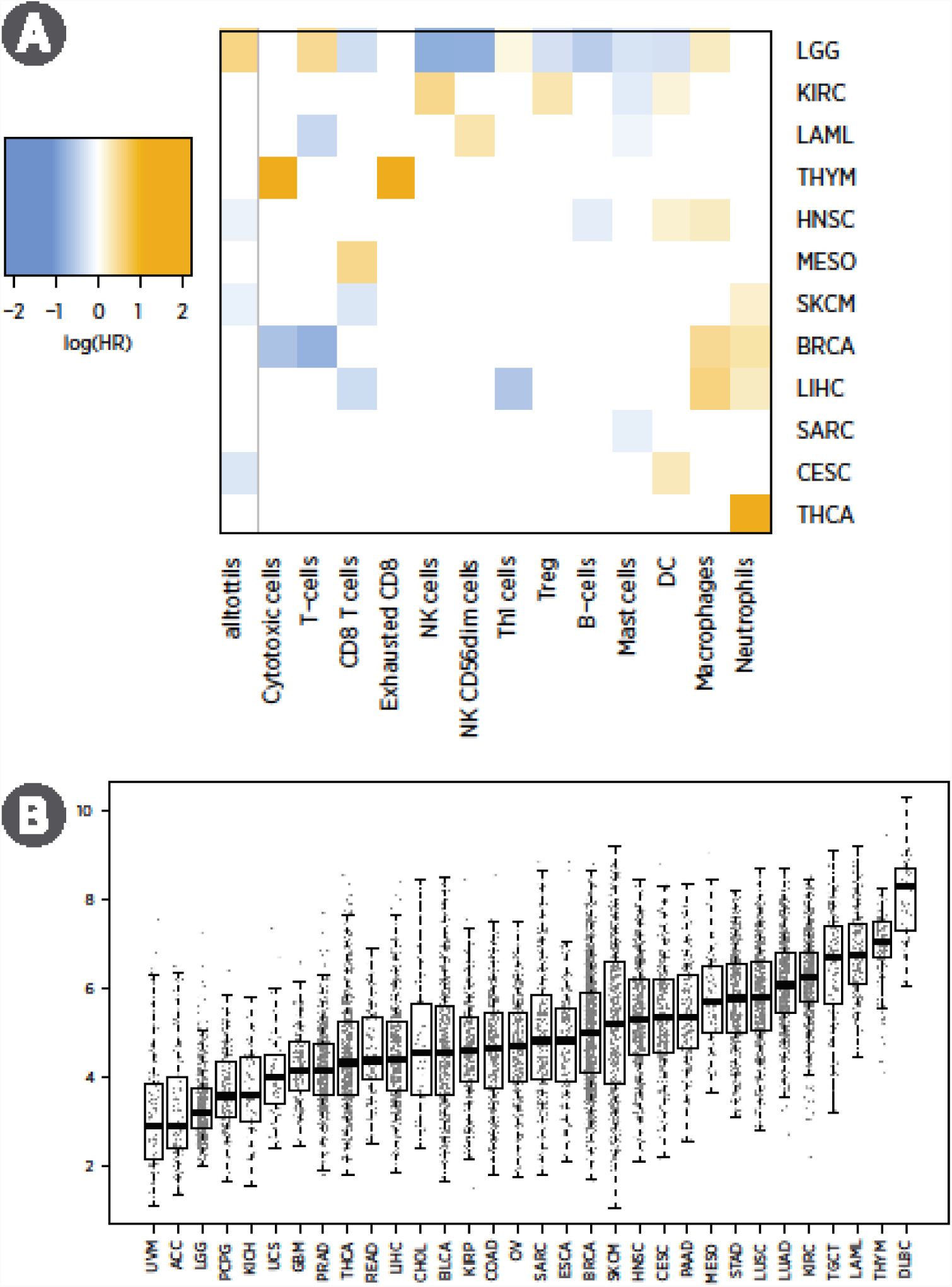
results of analyses of cell scores in TCGA RNASeq data. a) Prognostic information in cell scores. Orange indicates cell population enrichment within the total infiltrate is associated with poor outcome; blue indicates association with good prognosis. Only cells with at least one result with FDR < 0.05 are shown. b) Boxplot of total TIL abundance score across TCGA datasets.

#### Results in TCGA: prognostic significance of cell types

In each TCGA dataset, we tested the prognostic utility of total TILs and of each cell type’s enrichment or depletion relative to the total TILs. We first defined cell type enrichment scores as the residuals of linear regressions predicting each cell type from our total TILs score. These cell type enrichment scores do not measure absolute abundance of a cell type but rather its enrichment or depletion within the immune infiltrate. We then ran Cox regression predicting survival in each dataset from these cell type enrichment scores (Figure 6a.) Eleven tumor types had statistically significant (FDR < 0.1) [23] associations between survival and at least one feature of TIL abundance and makeup.

As others have shown [3,24], we see that immune cell populations have different prognostic implications in different tumor types, though some patterns are apparent. High total TILs score predicts longer survival in melanoma (SKCM) and head and neck (HNSC) tumors but worse prognosis in lower grade gliomas (LGG) and kidney renal clear cell tumors (KIRC). Enrichment of T-cells, CD8 T-cells and mast cells also tends to predict good prognosis. Enrichment of DCs, neutrophils and macrophages generally indicates poor prognosis, suggesting that these cell types mount a less effective immune response or can serve as suppressor cells.

The melanoma (SKCM) results best match the standard theory of immunotherapy: increased TILs and an infiltrate enriched for CD8 T-cells and Th1-induced IFN-gamma signaling indicate an effective immune response. The glioma (LGG) and kidney renal clear cell carcinoma (KIRC) results are striking: overall TIL abundance is associated with shorter survival, and most individual immune cell populations hold further prognostic importance. The LGG results can be explained by the danger of inflammation in the brain and by the role of macrophages in suppressive signaling. The thymoma (THYM) results are also interesting: although total TILs are not prognostic, there is rich prognostic information in the enrichment of various cell populations within the total infiltrate.

Of the 21 tumor types without evidence for a prognostic role of TILs, 10 lacked statistical power, with fewer than 33 events. The BLCA, CESC, COAD, ESCA, GBM, KIRP, LUAD, LUSC, OV, PAAD and STAD datasets all had at least 49 events, but lacked evidence for a prognostic role of TILs. The negative result in the colon cancer (COAD) dataset is a notable divergence from the prognostic relevance of the Immunoscore [2], though with only 49 events the dataset had modest power to establish an association.

The Supplementary Information contains further analyses of our cell scores in TCGA, including analyses of total immune abundance across tumor types, immune cell co-occurrence, correlation between immune populations and key immune oncology genes, relationships between tumor type and the makeup of the immune infiltrate, and associations between mutation burden and total immune infiltrate.

## Discussion

It is unknown whether our cell type scores track pure cell type abundance like flow and IHC or whether they track the product of cell type abundance and activity. For example, our data cannot rule out the possibility that highly active CD8 cells have increased expression of our CD8 marker genes relative to inactive CD8 cells. Whether cell type abundance or cell type activity levels have greater clinical relevance cannot be assessed by the data in this study.

CD4 subpopulations appear to lack sufficiently specific marker genes: for neither CD4 cells nor for any CD4 subpopulation did we find co-expression patterns among candidate genes consistent with cell type specificity. Quantification of these populations may require a deconvolution approach such as that employed by [9]. Alternatively, CD4 population functions could be measured with their canonical genes, similar to our use of TBX21 to measure Th1 cells and FOXP3 to measure Tregs.

Normalizing gene expression data for application of this method becomes complicated in non-tumor samples like PBMCs and cultured cells. In PBMCs, for example, it is likely that the standard reference genes have different expression levels in different cell types; thus a PBMC sample with abundant T-cells might be normalized to a different level than a PBMC sample with depleted T-cells. A workaround to this problem is to normalize not to reference genes but to a single immune cell population, yielding relative measurements of cell types like CD8 cells/CD3 cells and B-cells/CD3 cells. We apply this approach to our analysis of a flow cytometry dataset. Alternatively, normalizing to the average score of the major PBMC components – T-cells, B-cells, NK cells and macrophages – approximates normalization to the total number of cells in a sample.

To aid other investigators, we provide R code for calculating cell type abundances and for performing QC on marker genes in new datasets. We also list the candidate genes we examined (Table S3) and the validated genes (Table S4), and we provide cell type abundance scores on 9986 TCGA samples (Table S5). All data and code used in these analyses is provided in the Supplementary Information. NanoString Technologies has implemented this immune cell scoring method in a free, open-source analysis tool.

## Conclusions

We have identified a set of marker genes with sufficient cell type specificity that their expression levels can be used to measure immune cell subpopulations in the tumor microenvironment. Our method finds varying success across cell types (Figure 3d, Table 2), with some cell populations (T-cells, cytotoxic cells, mast cells, macrophages) having many well-behaved markers, with other cell populations possessing only weak markers (exhausted CD8 cells, NK CD56 dim cells), and others lacking any suitable markers (Th17 cells). Similarly, our 14 cell type gene lists have different levels of evidence. Our T cell and CD8 cell scores have the highest level of evidence: they correlate well with both IHC and flow, and their marker genes show strong marker-like behavior in TCGA data. We lack IHC data that could validate our B, NK, and NK CD56dim scores, but these cell scores correlate with flow cytometry and their markers behaved approximately as well in TCGA as our CD8 cell markers. Our mast cell, cytotoxic cell and macrophage scores have neither IHC nor flow measurements to support them, but their marker genes exhibited very strong marker-like co-expression in TCGA. Finally, our neutrophil and exhausted CD8 cell markers performed approximately as well in TCGA as our CD8 cell markers.

The immune cell scores described here can be implemented in a single assay using any gene expression platform, and any single cell type score can be calculated with an assay of just a handful of genes. Thus these cell scores represent a convenient technique for extracting detailed information about the tumor immune contexture. Furthermore, given their demonstrated prognostic value in TCGA and the increasingly well-understood associations between immune populations and response to immunotherapies [6], these cell scores may hold information useful for monitoring or predicting response to immunotherapy.

## List of abbreviations

IHC: immunohistochemistry
PBMCs: peripheral blood mononuclear cells
SD: standard deviation
RMSE: root mean squared error
TIL: tumor infiltrating leukocyte
NK: natural killer
DC: dendritic cell
CD: cluster of differentiation

## Declarations

### Ethics approval and consent to participate

PBMC and FFPE specimens of epithelial ovarian cancer patients were obtained with informed consent at Roswell Park Cancer Institute and University of Minnesota in accordance with approved protocols from the institutional review boards. Free and informed consent was obtained by Asterand for the FFPE samples used in the reproducibility section of the results.

### Author’s contributions

Patrick Danaher created the novel statistical method, helped identify candidate cell type marker genes, performed all analyses, and wrote the manuscript. Sarah Warren helped identify candidate cell type marker genes and contributed to the interpretation of results. Lucas Dennis helped define the aim of this research, helped identify candidate cell type marker genes and contributed to the interpretation of results. Leonard D’Amico ran and analyzed the flow cytometry experiments. Andrew White ran the NanoString experiments. Mary Disis helped acquire the tumor and PBMC samples used for this study, and she helped prepare the manuscript. Kunle Odunsi and Melissa Geller helped acquire the tumor and PBMC samples used for this study. Joseph Beechem helped define the aim of this research. Steven Fling helped run the trial from which our tumor and PBMC samples were derived, oversaw the generation of flow cytometry and IHC data, and helped prepare the manuscript.

## Acknowledgements

The results published here are in part based upon data generated by the TCGA Research Network: http://cancergenome.nih.gov/.

The authors would like to acknowledge the patients who provided tissue specimens to TCGA and to our own studies.

Martin McIntosh provided helpful insights that informed the method described here.

## References

1. Jochems C, Schlom J. Tumor-infiltrating immune cells and prognosis: the potential link between conventional cancer therapy and immunity. Experimental biology and medicine. 2011;236(5):567–79.

2. Galon J, et al. Cancer classification using the Immunoscore: a worldwide task force. Journal of translational medicine. 2012;205.

3. Fridman WH, Pagès F, Sautès-Fridman C, Galon J. The immune contexture in human tumours: impact on clinical outcome. Nature Reviews Cancer. 2012;12(4):298–306.

4. Salgado R, Denkert C, Demaria S, Sirtaine N, Klauschen F, Pruneri G, Wienert S, Van den Eynden G, Baehner FL, Penault-Llorca F, Perez EA. The evaluation of tumor-infiltrating lymphocytes (TILs) in breast cancer: recommendations by an International TILs Working Group 2014. Annals of Oncology. 2015;26(2):259–71.

5. Gentles AJ, Newman AM, Liu CL, Bratman SV, Feng W, Kim D, Nair VS, Xu Y, Khuong A, Hoang CD, Diehn M. The prognostic landscape of genes and infiltrating immune cells across human cancers. Nature medicine. 2015;21(8):938–45.

6. Topalian, SL., et al. Mechanism-driven biomarkers to guide immune checkpoint blockade in cancer therapy. Nature Reviews Cancer. 2016; 16(5):275–87.

7. Bindea G, et al. Spatiotemporal dynamics of intratumoral immune cells reveal the immune landscape in human cancer. Immunity. 2013;39(4):782–95.

8. Angelova M, Charoentong P, Hackl H, Fischer ML, Snajder R, Krogsdam AM, Waldner M, Bindea G, Mlecnik B, Galon J, Trajanoski Z. Characterization of the immunophenotypes and antigenomes of colorectal cancers reveals distinct tumor escape mechanisms and novel targets for immunotherapy. Genome Biol. 2015;16:64.

9. Newman AM., et al. Robust enumeration of cell subsets from tissue expression profiles. Nature methods. 2015;12(5):453–7.

10. Senbabaoglu Y, Winer AG, Gejman RS, Liu M, Luna A, Ostrovnaya I, Weinhold N, Lee W, Kaffenberger SD, Chen YB, Voss MH. The landscape of T cell infiltration in human cancer and its association with antigen presenting gene expression. bioRxiv. 2015;025908.

11. Vandesompele J, De Preter K, Pattyn F, Poppe B, Van Roy N, De Paepe A, Speleman F. Accurate normalization of real-time quantitative RT-PCR data by geometric averaging of multiple internal control genes. Genome biology. 2002;3(7):1–2.

12. Wherry EJ, Ha SJ, Kaech SM, Haining WN, Sarkar S, Kalia V, Subramaniam S, Blattman JN, Barber DL, Ahmed R. Molecular signature of CD8+ T cell exhaustion during chronic viral infection. Immunity. 2007;27(4):670–84.

13. Wherry EJ. T cell exhaustion. Nature immunology. 2011;12(6):492–9.

14. Gupta PK, Godec J, Wolski D, Adland E, Yates K, Pauken KE, Cosgrove C, Ledderose C, Junger WG, Robson SC, Wherry EJ. CD39 Expression Identifies Terminally Exhausted CD8+ T Cells. PLoS Pathog. 2015;11(10):e1005177.

15. Zhang M, Yao C, Guo Z, Zou J, Zhang L, Xiao H, Wang D, Yang D, Gong X, Zhu J, Li Y. Apparently low reproducibility of true differential expression discoveries in microarray studies. Bioinformatics. 2008;24(18):2057–63.

16. Galon, Jérôme, et al. “The continuum of cancer immunosurveillance: prognostic, predictive, and mechanistic signatures.” Immunity. 2013;39(1):11–26.

17. Sah NK, Khan Z, Khan GJ, Bisen PS. Structural, functional and therapeutic biology of survivin. Cancer letters. 2006;244(2):164–71.

18. Kim JM, Yamada M, Masai H. Functions of mammalian Cdc7 kinase in initiation/monitoring of DNA replication and development. Mutation Research/Fundamental and Molecular Mechanisms of Mutagenesis. 2003;532(1):29–40.

19. Ma L, Zhao X, Zhu X. Mitosin/CENP-F in mitosis, transcriptional control, and differentiation. Journal of biomedical science. 2006;13(2):205–13.

20. Bonilla FA, Fujita RM, Pivniouk VI, Chan AC, Geha RS. Adapter proteins SLP-76 and BLNK both are expressed by murine macrophages and are linked to signaling via Fcγ receptors I and II/III. Proceedings of the National Academy of Sciences. 2000;97(4):1725–30.

21. Barnett BE, et al. B Cell-Intrinsic T-bet Expression Is Required To Control Chronic Viral Infection. Journal of Immunology. 2016;pii: 1500368.

22. Hagemann AR, Hagemann IS, Cadungog M, Hwang WT, Patel P, Lal P, Hammond R, Gimotty PA, Chu CS, Rubin SC, Birrer MJ. Tissue-based immune monitoring II: multiple tumor sites reveal

23. Benjamini Y, Hochberg Y. Controlling the false discovery rate: a practical and powerful approach to multiple testing. Journal of the Royal Statistical Society. Series B (Methodological). 1995;289–300.

24. Linsley PS, Chaussabel D, Speake C. The Relationship of Immune Cell Signatures to Patient Survival Varies within and between Tumor Types. PloS One 10.9. 2015;e0138726.

